# The contribution of leaf sulfur content to the leaf economics spectrum explained by plant adaptive strategies

**DOI:** 10.1101/2020.12.16.423125

**Authors:** Michele Dalle Fratte, Simon Pierce, Magda Zanzottera, Bruno Enrico Leone Cerabolini

## Abstract

Sulfur is an essential macronutrient for plant metabolism. Terrestrial ecosystems have faced extensive anthropogenic sulfur depositions during the 20^th^ century, but currently decreasing trend of sulfur emissions suggest that it could become limiting, although its relationship with plant economics remains unclear.

We analysed leaf and nutrient traits for 740 vascular plant species growing in a wide range of environmental conditions. We aimed to determine whether leaf sulfur content (LSC) is associated with the leaf economics spectrum, and whether its distribution among functional types (life forms, leaf life span categories, and Grime’s CSR (Competitive, Stress-tolerant, Ruderal) strategies) could help to elucidate adaptive differences within plant taxa.

High LSC values corresponded mainly with R-, and to a lesser extent C-, strategy selection, hence the acquisitive extreme of plant economics. We found evidence of a relationship between nutrient stoichiometry and taxonomy, specifically at the acquisitive and conservative extremes of leaf economics. In general, LSC was significantly and positively correlated with leaf nitrogen content, but ruderal strategies in particular exhibited greater sulfur to nitrogen ratios.

Faced with a dearth of LSC data, leaf nitrogen content could thus be used as a coarse proxy of LSC within the context of plant economics. Different ratios among sulfur and nitrogen may be expected for ruderal species, suggesting that deeper insights from CSR strategies can provide a bridge between plant stoichiometry and ecology.

## 1. Introduction

Sulfur is one of the macronutrients essential for plant growth and physiological functions and is vital for metabolic processes (Aerts and Chapin 1999, Marschner 2012). Sulfur availability has a particularly important effect on photosynthesis because it is a component of the proteins involved in chlorophyll synthesis (Hu et al. 2017) and chloroplasts formation (Hasanuzzaman et al. 2018). Hence, sulfur limitation decreases yields and quality parameters of crops and the growth of tree species (Hawkesford and De Kok 2007). Demand for sulfur varies strongly between species (Buchner et al. 2004), but its uptake and assimilation are highly dependent on the availability of nutrients and on environmental conditions (Hawkesford and De Kok 2007). Sulfur assimilation is linked to multiple metabolic pathways responsible for a range of physiological functions, mainly including primary metabolite biosynthesis, stress responses, and pathogen defence (De Kok et al. 2017). Aside from effects on productivity, sulfur deficiency may thus also result in the loss of plant resistance to environmental stress and pests.

Together with nitrogen, sulfur is a component of essential organic compounds, such as proteins (amino acids cysteine and methionine), vitamins (biotin and thiamine), cofactors (Co-A and S-adenosyl methionine) and various other secondary compounds (Marschner 2012). Indeed, protein conformation (secondary and tertiary structure) depends on disulfide bridges between amino acid components, and sulfur is thus a fundamental requirement of living systems. Moreover, sulfur protects plants from biotic (i.e. herbivory; Badanes-Perez et al. 2014) and abiotic stresses such as salinity, drought, and toxic metals/metalloids (Hasanuzzaman et al. 2018), through different compounds that directly act as antioxidants or modulate antioxidant defence systems (e.g. glutathione, phytochelatins, metallothioneins) (Hawkesford and De Kok 2007). In addition, plants may emit minute amounts of volatile sulfur compounds, such as H_2_S, which may protect plants, for example against fungal attack (Hasanuzzaman et al. 2018).

Under natural conditions, the major sulfur source for plants are soil sulfates that originate from the mineralization of organic matter, which is sometimes a limiting step for plant sulfur supply, particularly in agricultural systems with low sulfur fertilisation. However, plants may acquire, through the leaves, other sources and forms of sulfur for growth, e.g. smaller quantities of sulfur dioxide (SO_2_; De Kok et al. 2017) coming from deposition following either natural (volcanic activity or sulfur springs) or anthropogenic (industrial) emissions (Bussotti et al. 2005). Indeed, until the 1980s, the amount of sulfur fertilization tended to increase along with SO_2_ emissions due to industrial activity, but simultaneously SO_2_ was the main contributor of the significant intensification of acid deposition such that sulfur was mainly regarded as a pollutant. Following the steady increase during the beginning of the 20^th^ century, national and international legislation on emission-reductions lead to the current worldwide trend of decreasing SO_2_ emissions (Aas et al. 2019), and to a sulfur deficiency scenario negatively affecting yield and quality of agricultural crops and biomass production of natural ecosystems. The extent to which sulfur could become a limiting element for both domesticated and wild species is unknown (Johnson et al. 2018). In order to understand and predict the effects of sulfur limitation, the resource economics of this nutrient (i.e. how plants manage and allocate mineral elements between different functions) must be determined. In other words, different species potentially exhibit different sulfur use in metabolism and concomitantly vary in growth responses, depending on life form and ecological strategy.

Despite the relevance of sulfur for plant metabolism, much attention has been paid to carbon, nitrogen and phosphorous, due to their larger contribution to plant growth, while few studies have investigated the stoichiometry of sulfur in terrestrial ecosystems (Han et al. 2011, Legay et al. 2014, Miatto and Batalha 2016, Shi et al. 2016, Wu et al. 2017), probably because it is a less critical limiting factor for plant productivity. Leaf nutrient contents of terrestrial plants are typically related to environmental factors (see Dalle Fratte et al. 2019a), but sulfur in particular can also be related to taxonomy (Zhang et al. 2012, Wu et al. 2017, De La Riva et al. 2018). For example, glucosinolates are sulfur-containing plant resistance compounds found mainly in species of the *Brassicaceae* family (Dijkshoorn and Van Wijk 1997), but have been recorded from 16 families in two main ‘mustard oil’ clades (Rodman et al. 1998). Among leaf nutrient contents, nitrogen has been identified as the main representative of global scale trade-offs between resource capture and conservation (Díaz et al. 2016). Such variation in plant resources allocation has been extensively investigated within the context of the leaf economics spectrum (LES; Wright et al. 2004, Reich 2014). Rapid resource acquisition is usually correlated with high values of specific leaf area (SLA) or leaf nitrogen content (LNC), while high leaf dry matter content (LDMC), lignin content or carbon to nitrogen ratio (C/N) reflect a resource conservation strategy (Pierce et al. 2007, Freschet et al. 2010, De La Riva et al. 2018). Since the larger portion of sulfur is used for primary metabolite biosynthesis, it is expected that leaf sulfur content (LSC) as well as the carbon to sulfur ratio (C/S) are embroiled in the LES.

The interactions of LSC with other plant traits has been studied only locally and/or for a few species (e.g. Bussotti et al. 2005, Sardans et al. 2008, Laliberté et al. 2012). De La Riva et al. (2018) highlighted that the use of functional groups (leaf life span and habitat type) appears to be useful to untangle the relationship between LSC and plant functioning. Grime’s CSR (Competitive, Stress-tolerant, Ruderal) adaptive strategies scheme (Grime 2006) defines functional groups identified on a solid theoretical base, and is the only ecological strategy theory that simultaneously explains both economics and size as fundamental gradients of plant adaptation and evolution (Grime and Pierce 2012). Considering that the extremes of the LES roughly correspond to stress-tolerant (conservative) and ruderal (acquisitive) CSR strategies (Díaz et al. 2016), these should also be connected to LSC. Despite this, we are not aware of comparative studies of LSC in relation to CSR strategies.

Here, based on a large dataset of the vascular flora of Northern Italy, representative of a wide range of environmental conditions, we tested whether: 1) LSC varies in concert with major axes of plant adaptation, in particular with the LES and consequently, 2) LSC distribution is related to specific life-forms, leaf life span categories and plant families along the LES, 3) LSC relates in different ways to CSR adaptive strategies so that these can help to explore LSC within taxonomic or functional groups.

## 2. Material and methods

### 2.1. Dataset

Our dataset (LIFTH, Leaf and nutrient Italian Flora Traits Hoard) consists of 740 species belonging to 99 families, of which only 17 are represented by at least 10 species records (Table A1). The dataset includes leaf and nutrient traits of both wild species characteristic of the main habitats of Northern Italy, and domesticated species (i.e. only cultivated in gardens and public parks). Nomenclature of every taxon (family, genus, species) in our dataset was standardized according to The Plant List (TPL, www.theplantlist.org) using the R package ‘Taxonstand’ (Cayuela et al. 2017).

For each species, within the same population we sampled from 5 to 15 fully expanded leaves selected randomly from the outer canopy of different individual adult plants growing in nature. All sampled populations were located far from possible anthropogenic SO_2_ emission sources, enough to suppose that there was no soil sulfur excess due to its depositions, also considering the high mobility of sulfate in soils (Johnson et al. 2018) in response to the negative trends of SO_2_ deposition observed over all European countries in the last two decades (EEA 2020). Sampling sites were widespread over an area of approximately 50,000 km^2^ ranging between 43°16’ − 46°34’ N latitudes and 07°54’ − 11°00’ E longitudes, with an altitudinal range of 2760 meters (30 - 2790 meters a.s.l.), i.e. from sea coasts to higher mountain belts. Indeed, even though most of the species in our dataset are representative of vegetation types of the Alpine (Southern Alps) and the Continental (Po Plain) Biogeographical Regions, an appreciable number of samples (7.3%) were collected within the Mediterranean region. Consequently, climate regimes, geological substrates and soils of the sampling sites show extensive variability, even at the local scale.

Laboratory measurements followed a standardized methodological protocol (Perez-Harguindeguy et al. 2016): we stored leaf material at 4°C overnight to obtain full turgidity for the determination of leaf fresh weight (LFW) and leaf area (LA; i.e. the surface area of fully expanded leaves); petioles and rachides were included as part of the leaf. LA was determined using a digital scanner and the software Leaf Area Measurement (LAM v.1.3; University of Sheffield, UK). Leaf dry weight (LDW) was then determined after drying for 24 h at 105°C, and parameters such as SLA (ratio between LA and LDW) and LDMC (ratio between LDW and LFW) were calculated. Hence, for each species we computed the mean value of all plant traits. We also derived canopy height (CANH) from literature data (Pignatti 1982). The dry leaf material was then mixed and pounded, and three randomly selected replicates were processed with a CHNS-analyzer (FlashEA 1112 series Thermo-Scientific), obtaining values of LCC (Leaf Carbon Content), LHC (Leaf Hydrogen Content), LNC (Leaf Nitrogen Content) and LSC. For each of the three replicates, we also calculated the ratio between LCC and LNC as well as LSC (respectively C/N and C/S), and averaged values for each species. Finally, using LA, SLA and LDMC, we calculated the C-, S-, and R-scores, according to the *StrateFy* tool of Pierce et al. (2017) and then classified species into seven categories including primary and secondary strategies (Grime, 2006): C (Competitive), CR (Competitive − Ruderal), R (Ruderal), SR (Stress-tolerant – Ruderal), S (Stress-tolerant), SC (Stress-tolerant – Competitive), CSR (C – S − R strategist). The *StrateFy* tool (Pierce et al. 2017) compares the trade-offs between LA, SLA and LDMC for each target species against worldwide variation in these traits, allowing the extent of acquisitive vs. conservative leaf economics and plant size to be quantified and compared against absolute global limits. This method mirrors variability in fourteen whole-plant, flowering, seed and leaf functional traits (Pierce et al. 2017) and has the advantage of using trait variation evident amongst vascular plants in general, including woody and herbaceous species and taxa such as ferns, and can thus be applied to large datasets of wild plant species and their communities.

We performed the analyses using nine plant traits: LA, LDW, SLA, LDMC, CANH, LNC, and C/N plus sulfur traits (LSC and C/S). We selected these traits because they represent the major axes of plant adaptation worldwide (Díaz et al. 2016) and/or are used for the computation of CSR scores (Pierce et al. 2017). We added LDW to account for the accumulation of dry matter of leaves aside from LA (Pierce et al. 2013), while C/S was included because it represents (along with C/N) a trait related to litter decomposition and carbon balance (Blair 1988, Freschet et al. 2010, Pierce et al. 2007).

We categorised species into four life form categories following Dalle Fratte et al. (2019b): trees (n = 86), shrubs (n = 118), long-lived herbs (n = 433), and short-lived herbs (n = 103). Such broad categories largely correspond to those of Raunkiær (1934) and help to remove growth forms composed of few species (e.g. lianas, herbaceous vines, rushes). Essentially, this classification divides woody species (n = 204) into trees and shrubs (shrubs and lianas), and non-woody species (n = 536) on the base of their life cycle length, regardless of being graminoids, forbs, aquatics or herbaceous vines: short-lived (which include annual and biennal) vs. long-lived (perennial) herbs.

We also classified species according to their leaf life span, which plays a critical role in plant economics (Reich 2014). We adopted three categories: evergreen (n = 64), wintergreen (n = 308), deciduous (n = 368). Accordingly, evergreen are those plants having leaves or needles all year round and older than one year, while deciduous have leaves that die off after having been green during summer, and only rarely some leaves may remain green during mild winters. Wintergreen species are those plants that build leaves during the vegetation period, but they remain green until the next leaf unfolding in spring unless they are exposed to very low temperatures or extreme drought.

### 2.2. Statistical analysis

We computed all the statistical analyses using the R software (R Core Team 2020). Data were first normality checked by means of the Shapiro-Wilk test and accordingly we transformed all variables by logarithmic function [*log*(x)], except LDMC for which we used the square root function [*sqrt*(x)].

To highlight the main adaptive trends within the trait space of our dataset and to seek relationships of sulfur traits (LSC, C/S) with these, we first performed a principal component analysis (PCA), centred by standard deviation and followed by varimax rotation, using the function ‘principal’ of the package ‘psych’ (Revelle 2018). The number of significant dimensions were identified by comparing the eigenvalues of each component with the value given by the broken stick distribution, and statistically significant correlations between PCA axes and plant traits were identified using Pearson’s correlation coefficient.

We applied one-way analysis of variance (ANOVA) with Tukey post-hoc comparisons to find differences of plant traits among the categories of life form, leaf life span and CSR plant strategies. In order to exclude influences of phylogenetic correlation we used linear mixed-effects models considering the family as a random effect. For each ANOVA, we tested the effect of family comparing the results of the linear mixed-effects model with those of a general linear model without any random effect (Dalle Fratte et al. 2019a). The addition of family on the models was always highly significant (p < 0.01). We also applied simple one-way ANOVA with Tukey post-hoc comparisons to test differences of plant traits among plant families, considering only those represented by at least 10 species.

We used simple linear regression models to evaluate the log-log relationship of LSC (dependent variable) with LNC (independent variables) within the whole dataset, as well as within each single category of life form, leaf life span, CSR plant strategies, and within each plant family (represented by at least 10 species). The log-log regression has been evaluated as a robust method for estimating nutrient ratios (Isles 2020), and has been showed to provide effective insights concerning nitrogen and sulfur acquisition (Legay et al. 2014; Wu et al. 2017). We considered the R^2^ to assess the amount of variation explained by each regression model, and to identify the relation of the ratio between LSC and LNC among plant families and CSR plant strategies, we correlated the slope of the linear regression (LSC vs. LNC) with the average values of family’s CSR scores. We used the package ‘nlme’ (Pinheiro et al. 2018) for the linear mixed-effects model, the base R package ‘stats’ for general linear model and simple linear regression, and the package ‘ggtern’ (Hamilton and Ferry 2018) for the ternary visualization of CSR strategies.

## 3. Results

The main principal adaptive trends of species in the PCA were represented by a two-dimensional space (cumulative percentage of variance = 69%, Fig. A2) defined by variation in the LES (PC1 = 45%) and of the size and the dimensions of leaves and plant height (PC2 = 24%; Fig. 1). The first axis (PC1) correlated positively with SLA and LNC, and negatively with LDMC and C/N (Table 1), indicating variation in trait values ranging from acquisitive strategies to values indicating conservative resource use, while PC2 correlated positively with CANH, LDW and LA (i.e. towards taller plants with larger and heavier leaves). Sulfur traits (LSC and C/S) correlated significantly with PC1 and grouped, respectively, with acquisitive and conservative traits.

**Table 1:** Pearson’s correlation coefficients (r) of the first two principal component analysis (PCA) axes with traits values for the 740 plant species selected for the analysis. Emboldened values are highly correlated (r > ± 0.5) and statistically significant at p ≤ 0.01 level (critical value of r = 0.09, d.f. = 739). Legend: LA = leaf area, LDW = leaf dry weight, CANH = canopy height, SLA = specific leaf area, LNC = leaf nitrogen content, LSC = leaf sulfur content, LDMC = leaf dry matter content, C/N = carbon to nitrogen ratio, C/S = carbon to sulfur ratio.

|  | **PC1** | **PC2** |
| --- | --- | --- |
| LA | 0.25 | **0.90** |
| LDW | 0.05 | **0.95** |
| CANH | -0.31 | **0.67** |
| SLA | **0.73** | -0.24 |
| LNC | **0.85** | 0.17 |
| LSC | **0.79** | 0.05 |
| LDMC | **-0.70** | 0.20 |
| CN | **-0.89** | -0.14 |
| CS | **-0.81** | -0.04 |

**Figure 1:**
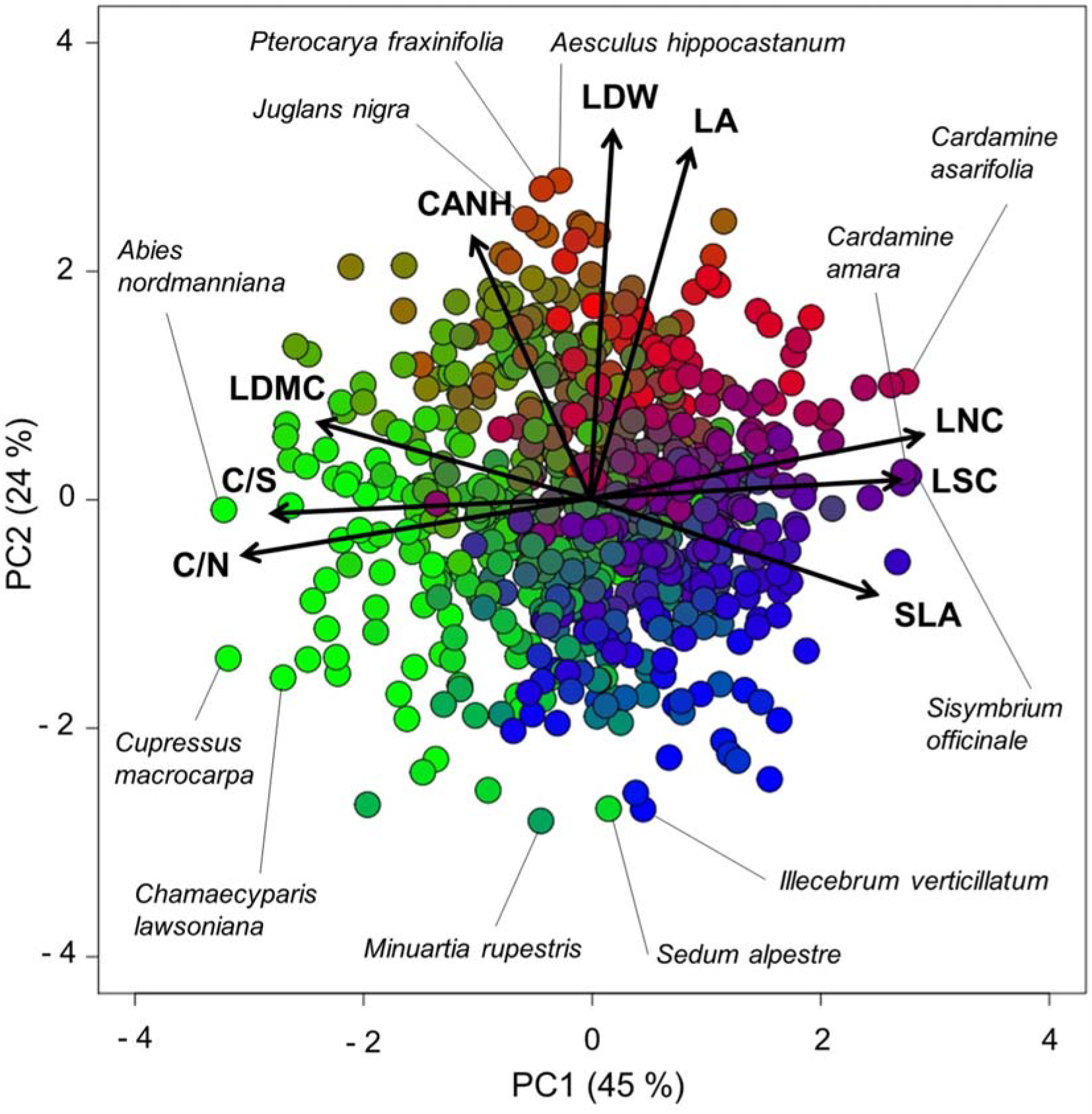
Principal Component Analysis on traits values of the 740 vascular plant species selected for the analysis. Species points are coloured by RGB classification according to their CSR scores calculated with *StrateFy* (Pierce et al. 2017): red = C, green = S, blue = R. Labelled points are species with the three highest and lowest values on PC1 and PC2. Legend: CANH = canopy height, C/N = carbon to nitrogen ratio, C/S = carbon to sulfur ratio, LA = leaf area, LDMC = leaf dry matter content, LDW = leaf dry weight, LNC = leaf nitrogen content, LSC = leaf sulfur content, SLA = specific leaf area.

The LES was thus well defined by six traits, including LSC and C/S, which also showed statistically significant differences among life form categories (Fig. 2). Trees and shrubs exhibited the lowest mean values of all the acquisitive traits (SLA, LNC and LSC), while short-lived herbs displayed the highest mean values of SLA and LSC. Only for LNC, short and long-lived herbs did not exhibit significant differences among each other. With regard to the set of conservative traits, trees and shrubs showed the highest mean values of C/N and C/S, while trees displayed significantly higher values of LDMC compared to shrubs. Short-lived herbs were the less conservative, showing the lowest LDMC and C/S, but similar values of C/N compared to long-lived herbs. With regard to leaf life span categories, evergreen species were more conservative compared to deciduous and wintergreen, which were indeed more acquisitive.

**Figure 2:**
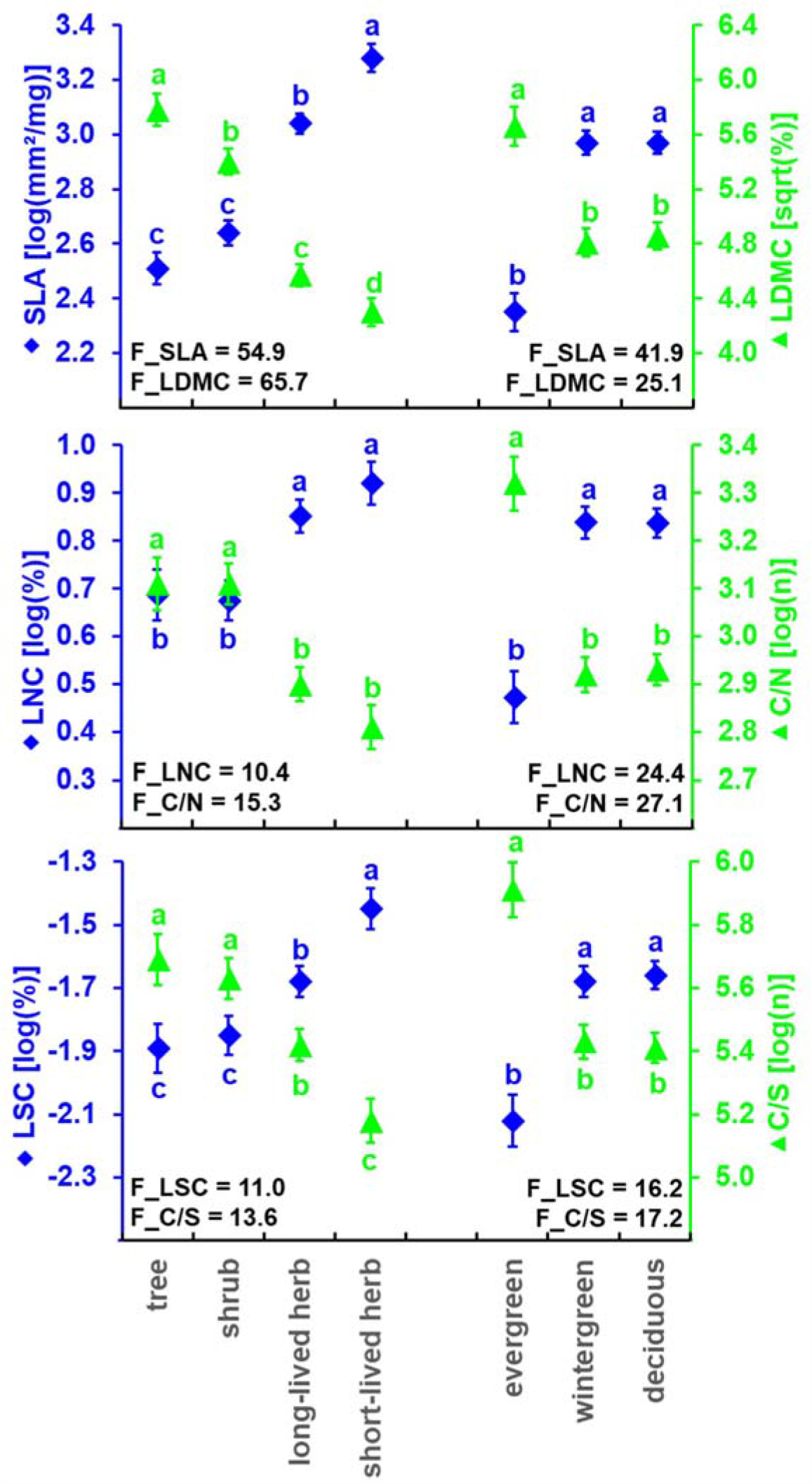
Mean values and standard error (vertical lines) of traits involved in the LES (Leaf Economics Spectrum) for each life form and leaf life span category. Small letters represent post-hoc comparisons. All the ANOVAs were highly significant (p < 0.001, F-value reported in the figure). Legend: C/N = carbon to nitrogen ratio, C/S = carbon to sulfur ratio, LDMC = leaf dry matter content, LNC = leaf nitrogen content, LSC = leaf sulfur content, SLA = specific leaf area.

Only a few plant families exhibited significant differences of the mean values of traits correlated with the LES (Fig. A3). With regard to leaf traits, such as SLA and LDMC, the *Pinaceae* family was found to be the most conservative, while *Orchidaceae, Brassicaceae* and *Caryophyllaceae* were the most acquisitive, despite not showing significant differences compared to many families with intermediate characteristics. Nutrient traits confirmed *Pinaceae* and *Brassicaceae* to be families that were significantly different from others, respectively placed at the conservative and acquisitive extremes of the LES. Nitrogen and sulfur traits showed a similar pattern, even though LNC and C/N discriminated more the *Pinaceae*, while LSC and C/S the *Brassicaceae*.

Ruderal (R) species showed on average the highest LSC values, similar only to competitive (C) species or species with both competitive and ruderal characteristics (CR), in contrast to S-selected species, which showed the lowest values, only comparable to those of the SC category (Fig. 3b). Only a few species, among those with higher LSC, were placed toward the centre of the CSR triangle (*Erysimum rhaeticum* and *Mercurialis annua*) or even toward the S-selected corner (*Triglochin palustris, Equisetum fluviatile, E. variegatum*) (Fig. 3a).

**Figure 3:**
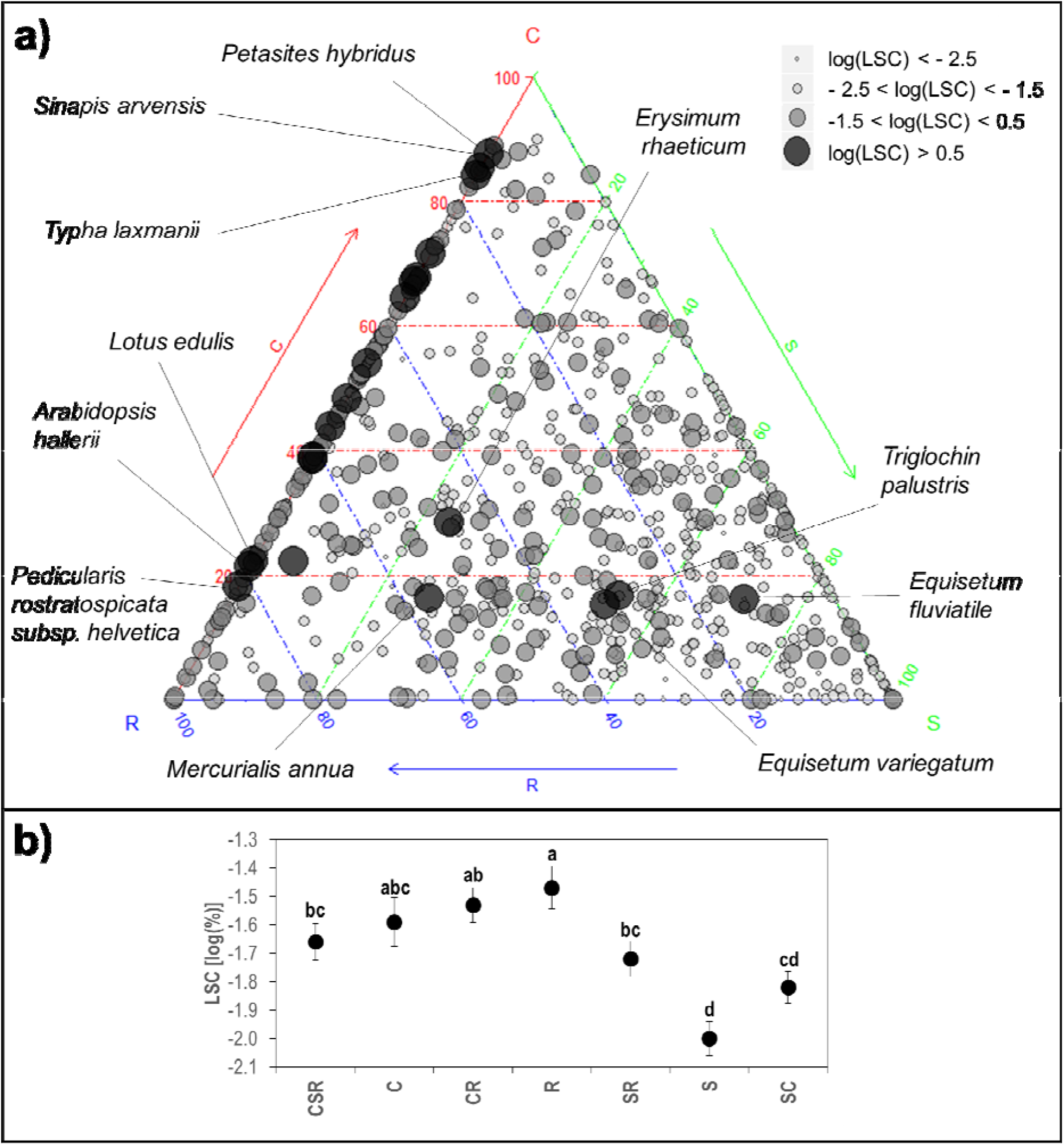
(a) Ternary visualization of CSR (Competitive, Stress-tolerant, Ruderal) strategies of the 740 species selected for the analysis; darker and wider points denote higher values of LSC (Leaf Sulfur Content). (b) Mean values and standard error (vertical lines) of LSC among plant strategies categories (F = 11.8, p < 0.0001). Small letters represent post-hoc comparison.

LSC showed a significant positive linear relationship with LNC (Table 2) considering all species together (slope = 0.81, R^2^ = 0.30, p < 0.01), which was even more robust within the R strategy category (slope = 0.95, R^2^ = 0.42, p < 0.01). With regard to families, we found an increase of both slope and R^2^ compared to the overall equation for *Orchidaceae, Poaceae, Juncaceae, Rosaceae*, and *Caryophyllaceae*. Finally, we observed a positive correlation between the mean R scores and the slope values of the linear regression equations LSC vs. LNC calculated for each plant family (Fig. 4). *Caryophyllaceae* and *Orchidaceae* were the two families with the highest mean R scores and consequently showed the highest slope values (LSC vs. LNC), while *Apiaceae* and *Salicaceae* were found at the opposite extreme of this gradient.

**Table 2:** Coefficients of simple linear regressions between LSC (dependent variable) and LNC (independent variable) for all species and different categories of life form, leaf life span, Grime’s CSR strategies, as well as for those families represented by at least 10 species in the dataset. Emboldened values are those with R^2^ higher than the one of the overall equation. Legend: n = number of species, α = intercept, β = slope, p = p-value (ns = not significant, * = p < 0.05, ** = p < 0.01, *** = p < 0.001).

| | | n | $\alpha$ | $\beta$ | $R^2$ | p |
| --- | --- | --- | --- | --- | --- | --- |
| **ALL SPECIES** |  | 740 | -2.37 | 0.81 | 0.30 | *** |
| **LIFE FORM** | **tree** | 86 | -2.36 | 0.56 | 0.21 | *** |
|  | **shrub** | 118 | -2.35 | 0.71 | 0.25 | *** |
|  | **long-lived herb** | 433 | -2.38 | 0.82 | 0.29 | *** |
|  | **short-lived herb** | 103 | -2.12 | 0.73 | 0.22 | *** |
| **LEAF LIFE SPAN** | **deciduous** | 368 | -2.22 | 0.63 | 0.19 | *** |
|  | **wintergreen** | **308** | **-2.43** | **0.93** | **0.32** | *** |
|  | **evergreen** | 64 | -2.46 | 0.59 | 0.15 | ** |
| **CSR STRATEGIES** | **C** | 44 | -2.29 | 0.84 | 0.20 | ** |
|  | **CR** | 128 | -2.43 | 0.90 | 0.23 | *** |
|  | **CSR** | 93 | -2.20 | 0.60 | 0.12 | *** |
|  | **R** | **69** | **-2.41** | **0.95** | **0.42** | *** |
|  | **SR** | 119 | -2.36 | 0.84 | 0.21 | *** |
|  | **S** | 144 | -2.36 | 0.68 | 0.22 | *** |
|  | **SC** | 143 | -2.25 | 0.57 | 0.14 | *** |
| **FAMILY** | *Pinaceae* | 17 | -2.54 | 0.97 | 0.25 | * |
|  | *Asparagaceae* | **12** | **-2.40** | **0.75** | **0.55** | ** |
|  | *Orchidaceae* | **13** | **-3.48** | **1.54** | **0.52** | ** |
|  | *Poaceae* | **75** | **-2.51** | **1.08** | **0.54** | *** |
|  | *Juncaceae* | **16** | **-2.29** | **1.01** | **0.40** | ** |
|  | *Cyperaceae* | 70 | -2.21 | 0.59 | 0.17 | *** |
|  | *Salicaceae* | 13 | -1.58 | 0.20 | 0.01 | ns |
|  | *Ranunculaceae* | 13 | -2.15 | 0.54 | 0.10 | ns |
|  | *Leguminosae* | 39 | -2.75 | 0.85 | 0.21 | ** |
|  | *Rosaceae* | **40** | **-2.78** | **0.90** | **0.46** | *** |
|  | *Brassicaceae* | 19 | -1.74 | 0.86 | 0.20 | * |
|  | *Caprifoliaceae* | **13** | **-2.28** | **0.69** | **0.38** | * |
|  | *Caryophyllaceae* | **21** | **-2.81** | **1.18** | **0.64** | *** |
|  | *Lamiaceae* | 13 | -2.05 | 0.49 | 0.18 | ns |
|  | *Plantaginaceae* | 16 | -1.94 | 0.54 | 0.17 | ns |
|  | *Compositae* | 66 | -2.32 | 0.90 | 0.21 | *** |
|  | *Apiaceae* | 23 | -1.63 | 0.06 | 0.00 | ns |

**Figure 4:**
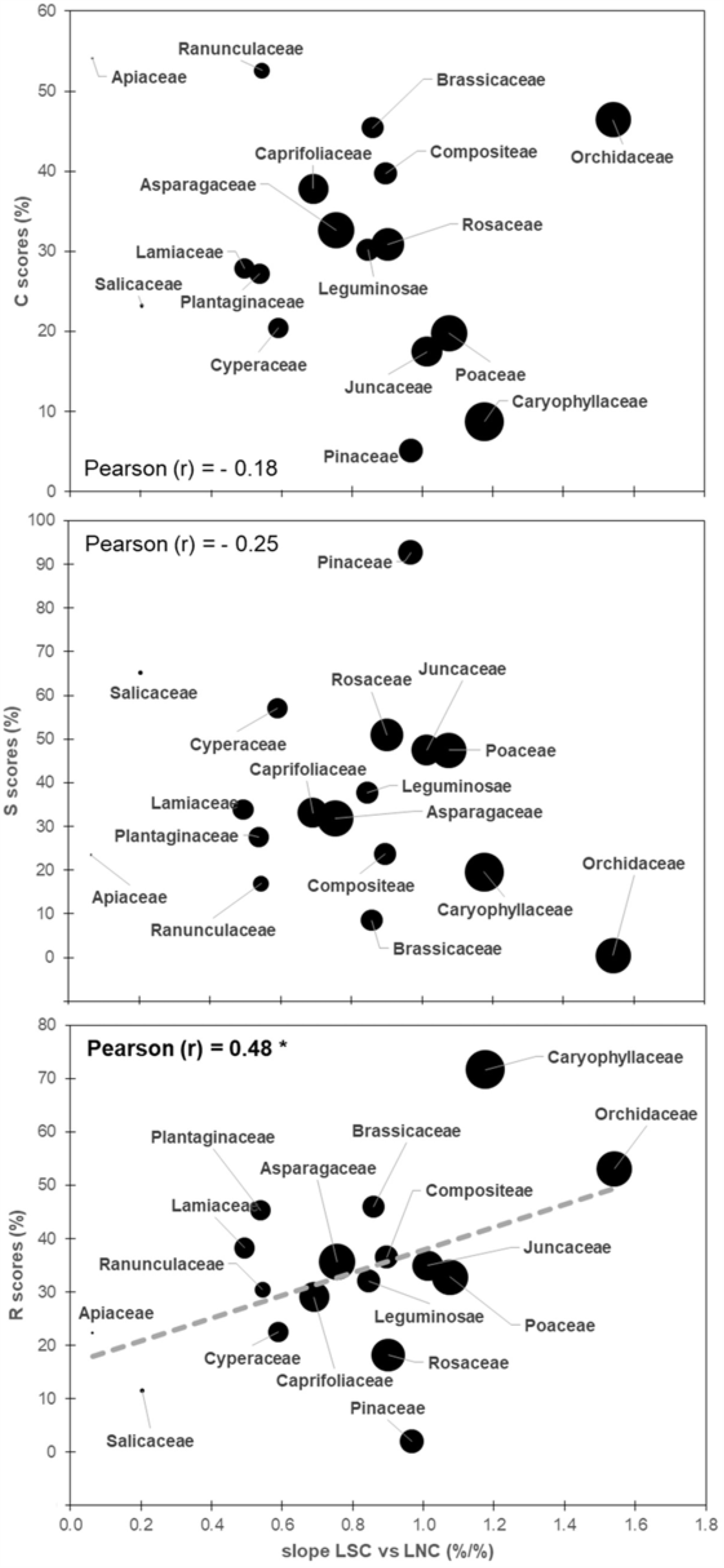
Relationship between slope values resulted from the linear regression LSC vs. LNC within plant families represented by at least 10 species, and their mean values of C-, S- and R-selection scores. Larger points denote higher R^2^ values of the linear regression LSC vs. LNC (see Table 2). Pearson’s correlation coefficients (r) is reported in each figure; the emboldened one is significant at p ≤ 0.05 (critical value = 0.48, d.f. = 15).

## 4. Discussion

Firstly, our analysis at the regional scale confirmed the robustness of the two main axes of variation of plant functioning: representing the LES (i.e. fast-slow leaf economics; Reich 2014), and the size and dimensions of plant and leaves (both comprising the global spectrum of plant form and function; Díaz et al. 2016; Fig. 1 and Table 1). The pattern of SLA, LNC, together with LSC at one extreme and of LDMC, C/N, together with C/S at the other, reflected, respectively, the acquisitive-to-conservative trade-off determining the LES (Wright et al. 2004, Reich 2014), demonstrating that variation in LSC was associated with variation in the LES, supporting Hypothesis 1. Indeed, in our study the LES was the dimension accounting for the largest source of variance. The main reason for this apparent discrepancy with respect to the global spectrum may be related to the lower sample size but also to the different set of traits used in our analyses and, specifically, the inclusion of sulfur traits. This leads to the principal finding of our work, namely that LSC and C/S are in strong agreement with variation in the traits normally used for global spectrum studies, given their evident contribution to the LES axis. This also suggests that additional nutrients other than carbon, nitrogen and phosphorous, less frequently included in functional trait analyses, may also be embroiled in the LES (Sardans et al. 2008, Laliberté et al. 2012, De La Riva et al. 2018). Plant growth requires at least 17 mineral elements used in leaves to support fundamental physiological processes (Marschner 2012). Accordingly, SLA and leaf nutrient contents are supposedly closely related across species (Niinemets and Kull 2003, Wright and Cannon 2001, Wright et al. 2004).

We observed a significant linear relationship between LSC and LNC that confirmed previous similar results (Legay et al. 2014, Wu et al. 2017, Dijkshoorn and Van Wijk 1997; Table 2), highlighting that the assimilation processes of sulfur and nitrogen are associated (Hasanuzzaman et al. 2018), as most available reduced nitrogen and sulfur are incorporated into amino acids and subsequently into proteins (De Kok et al. 2017). Nitrogen together with phosphorous are integral to proteins of the photosynthetic apparatus, including Rubisco, and their contents are thus positively correlated with net CO_2_ assimilation rate, dark respiration rate and relative growth rate (Wright et al. 2004, Reich et al. 2008). In addition to chloroplasts formation (Scherer 2008), sulfur has specific roles in fundamental processes, from photosynthesis to carbon and nitrogen metabolism (Droux 2004), including roles not shared by nitrogen and phosphorous but relevant to key biochemical pathways: for example, it contributes to ferredoxin oxidation (De Kok et al. 2017), and is part of Iron-sulfur clusters that aid the production of protein sufB, required for chlorophyll production (Hu et al. 2017). Sulfur thus represents an additional limitation to chlorophyll content and photosynthetic capacity with respect to nitrogen contents (Terry 1976, Resurreccion et al. 2001), and is vital to the achievement of the higher photosynthetic and respiration rates that characterize acquisitive species (Wright et al. 2004).

In contrast, conservative species exhibit greater mechanical support (e.g. high LDMC) and low photosynthetic and respiration rates, which involves greater leaf construction costs and times (Wright and Cannon 2001, Wright et al. 2004). C/N and C/S provide an indication of the relative investment in structure (carbon) and cell functioning (nitrogen and sulfur) and thus represent alternative measures of resource economics with respect to SLA and LDMC (Freschet et al. 2010), which use just mass measurements, also of clear physiological relevance. Indeed, physiological rates such as photosynthetic capacity and respiration rate are “carbon traits” (as opposed to “water or nutrient traits”; Reich 2014) that determine the amount of organic (carbon-based) matter a plant can produce. Species having leaves with a higher C/N are usually slow growing (Pierce et al. 2007, Freschet et al. 2010) with higher lignin content, and this optimization has also been found to relate to defence against herbivory (Hanley et al. 2007, Moreira et al. 2018). Thus, the addition of C/S to this framework could aid predictions concerning plant resistance to herbivory, as it expresses not only leaf digestibility (which corresponds with high leaf nutrient contents) but it also informs with regard to plant toxicity provided by sulfur-based plant defence metabolites (e.g. Badanes-Perez et al. 2014).

Leaf digestibility can subsequently alter litter decomposability (Cornelissen et al. 2004). Indeed, C/N has often been used as a proxy of environmental variables such as soil organic matter quality and litter decomposition rates (Freschet et al. 2010). As a rule, carbon-to-nutrient ratios are important determinants of whether an element will be immobilized or released as litter decomposition proceeds (Blair 1988). Below a critical threshold, nutrients will be incorporated in soil microbial biomass and by-products as carbon are mineralized, thus lowering carbon-to-nutrient ratios. Plant community composition and soil fertility interact through complex feedbacks (Aerts and Chapin 1999) that are detachable by traits related to the LES (Freschet et al. 2010). Based on our findings, it is possible to extend the same interpretation of C/N, in terms of litter decomposability, also to C/S. Accordingly, LSC could influence litter decomposition rates by altering the structure of microbial communities as well as fungi development (Legay et al. 2014). LSC has therefore a significant connection to soil carbon dynamics (Miatto and Batalha 2016, Shi et al. 2016) and to nutrient economy at the community level (Laliberté et al. 2012), which was recently found to reflect the variation from communities with resource-acquisitive (high leaf nitrogen and phosphorous content) to those with resource-conservative characteristics (high LCC) (Bruelheide et al. 2018).

Leaf structure is considered one of the main drivers of the LES (Wright et al. 2004) and plant functional types have often been invoked to explain differences existing along the LES (Reich 2014, Díaz et al. 2016). In this context, we detected significant differences classifying species into broad functional types, that is, within both life form and leaf life span categories. Trees and shrubs, as well as evergreen species, showed higher affinities for conservative strategies, in contrast to short and long-lived herbs, as well as wintergreen and deciduous species, which overall demonstrated relatively acquisitive strategies. This interpretation was supported by differences in mean values of all analysed leaf traits, including sulfur traits (Fig. 2). Evergreen species, compared to wintergreen and deciduous, require greater mechanical support, so that the increase of LDMC and the corresponding decrease of SLA are related to a greater portion of carbon in structural tissue (Villar and Merino 2001), which makes them less susceptible to environmental hazards and stress (Bussotti et al. 2005, Sardans et al. 2008, Poorter et al. 2009). They also displayed lower LNC and LSC, associated with higher C/N and C/S, indicating low nutrient requirements due to a more efficient use, typical of slow growing species (De La Riva et al. 2018), despite the greater energetic costs of tissue construction (Villar and Merino 2001).

We observed similar large differences in traits related to the LES among woody and non-woody life forms, confirming that herbs, particularly short-lived herbs, are more acquisitive than trees and shrubs. Our results show once more that the use of life forms can help to discriminate variations along the gradients underlying the LES (Wright et al. 2005) and nutrient use efficiency, as herbaceous species, particularly short-lived ones, are those with higher leaf nutrient contents (Han et al. 2011), sulfur included. Nevertheless, at the global scale woody and herbaceous species have shown extensive overlap along the LES (Díaz et al. 2016), partially due to the larger number of species and range of climates that were considered in the global study. However, we found only herbaceous species at the most acquisitive extreme of the LES, confirming their adaptation to include relatively acquisitive traits (Pierce et al. 2013), as opposed to trees, which instead have a more conservative set of traits.

We found evidence that high levels of LSC strongly relate to the ruderal strategy (the extent of R-selection) and, to a lesser extent, to the competitive strategy (C-selection), while the stress tolerant strategy (S-selection) was characterized by the lowest values, confirming the predictive strength of Grime’s CSR adaptive strategies also regarding nutrient economics (Pierce et al. 2007) (Fig. 3). Only a few species with a high LSC were located towards the stress-tolerant (S-selection) corner; specifically, two species of the genus *Equisetum* (*E. fluviatile* and *E. variegatum*). This is not surprising if we consider that *Equisetaceae* are known to have high nutrient contents (Marsh et al. 2000) due to an efficient nutrient uptake that allows these species to thrive under a wide range of conditions (Husby 2013). Moreover, they exhibit an abnormally high accumulation of silica, up to 25% of dry weight (Gierlinger et al. 2008) that, combined with the development of photosynthetic stems, may provide heavier structures and hence apparent affinities to stress-tolerant strategies. Among all the CSR strategies categories, ruderal species showed the closest linear relationship between LSC and LNC (R^2^ = 0.42) and the steepest slope (β = 0.95), i.e. the greatest portion of sulfur compared to nitrogen (Table 2). This led us to hypothesize that the ratio of the two nutrients may be more stable within this subset of species, which must invest in a rapid growth to complete faster the lifecycle, in order to maintain the population in the face of disturbances events (Grime 2006, Grime and Pierce 2012, Pierce et al. 2017).

Leaf nutrient contents are also linked to taxonomy (Zhang et al. 2012, Miatto and Batalha 2016), as we clearly observed for the *Pinaceae* (and other conifers) and for the *Brassicaceae*, respectively placed at the conservative and acquisitive extremes of the LES (Fig. 1 and Fig. A3). *Brassicaceae* displayed the highest values of LSC, probably because of the production of sulfur-containing glucosinolates (Dijkshoorn and Van Wijk 1997, Badanes-Perez et al. 2014). They also showed high values of LNC, which was very generally but significantly correlated with LSC (R^2^ = 0.20, p < 0.05; Table 2), perhaps because both nutrients are employed to various extents throughout primary and secondary metabolism. A tighter linear relationship between LSC and LNC (i.e. higher values of R^2^; Table 2) was instead shown by other families with a higher degree of ruderality than *Brassicaceae*, like *Orchidaceae* and *Caryophyllaceae*, which also showed a higher LSC to LNC ratio (i.e. higher values of the slope β; Table 2). This evidence strengthens the overall linear relationship found for ruderal strategies, but also suggests that, considering families, the greater it is the extent of R-selection, the more the portion of sulfur compared to nitrogen in leaves is high and stable (Fig. 4). Explaining high values of LSC, and their constancy in relation to LNC, is beyond the scope of the present study, and likely involve many other products of the secondary metabolism involving sulfur compounds (Hawkesford and De Kok 2007).

The biological stoichiometry of plants can play a key role in exploring evolutionary processes and adaptive variation in the biota, thus is crucial in evaluating ecological responses to global change (Elser et al. 2010). We observed high relevance of sulfur for plant functioning that can be integrated with other plant traits to learn more about the processes and patterns of ecosystem development in response to environmental changes. Future changes of nutrient deposition loads, together with climate change, will determine new environmental scenarios that may substantially alter the chemical composition of terrestrial ecosystems, with profound consequences for competition among species, plant community composition and biogeochemical cycles (Shi et al. 2016). Specifically, the biogeochemical cycle of sulfur is (and has been) largely impacted by anthropogenic activities so that it is necessary to understand its contribution within plant functioning and how it could affect the kind of species (i.e. with fast or slow economics) that can grow in a given environment. However, few LSC data appear to be available today in international trait databases (e.g. in TRY, Kattge et al. 2020), compared to other leaf nutrient contents, particularly LNC. In light of this, here we presented a large dataset of leaf traits, including LNC and LSC as relevant, covering a wide range of plant forms and functions, useful to integrate, at least at the regional scale, the understanding of plant functional responses to global change.

## 5. Conclusions

The link identified here between sulfur traits, LSC and C/S, and other leaf traits underlying the LES provides insights concerning the role of sulfur for plant functioning. The interaction between sulfur and other nutrients, specifically LNC and C/N, suggests that sulfur traits also scale up to ecosystem properties, such as biomass production or litter decomposability, which are strictly related with LES traits. We also observed a positive correlation between LSC and LNC, which showed different patterns throughout broad functional type categories. Among these, Grime’s CSR adaptive strategies were found to be the most reliable, demonstrating their wide applicability also in connecting knowledge of sulfur physiology to plant ecology. Specifically, we found that species with high LSC were associated with the acquisitive extreme of the LES detected at the global scale, which was specifically represented by highly R-selected (ruderal) strategies.

## Acknowledgements

We thank Andrea Gianotti, Matteo Francocci and Martina Guglielmi for assistance in the field and in the laboratory.

## Data availability statement

On acceptance, all data used for these analyses will be deposited at TRY Plant Trait Database https://try-db.org/TryWeb/Home.php

## Author contribution

**Michele Dalle Fratte**: Conceptualization, Methodology, Investigation, Visualization, Writing-Original draft preparation, Writing-Reviewing and Editing. **Simon Pierce**: Writing-Reviewing and Editing. **Magda Zanzottera**: Investigation, Writing-Original draft preparation, Writing-Reviewing and Editing. **Bruno E**.**L. Cerabolini**: Conceptualization, Methodology, Writing-Reviewing and Editing, Supervision.

## Appendices

### Appendix 1 (Table A1)

List of all plant families considered in the analysis (number of records = 740). Emboldened families are those represented by at least 10 species, selected for the regression analysis between LSC (Leaf Sulfur Content) and LNC (Leaf Nitrogen Content).

| **Family** | **n°** | **Family** | **n°** | **Family** | **n°** |
| --- | --- | --- | --- | --- | --- |
| **Poaceae** | **75** | Aspleniaceae | 5 | Lycopodiaceae | 2 |
| **Cyperaceae** | **70** | Boraginaceae | 5 | Moraceae | 2 |
| **Compositae** | **66** | Ulmaceae | 5 | Oxalidaceae | 2 |
| **Rosaceae** | **40** | Violaceae | 5 | Pteridaceae | 2 |
| **Leguminosae** | **39** | Equisetaceae | 4 | Rhamnaceae | 2 |
| **Apiaceae** | **23** | Liliaceae | 4 | Santalaceae | 2 |
| **Caryophyllaceae** | **21** | Magnoliaceae | 4 | Taxaceae | 2 |
| **Brassicaceae** | **19** | Papaveraceae | 4 | Actinidiaceae | 1 |
| **Pinaceae** | **17** | Primulaceae | 4 | Altingiaceae | 1 |
| **Juncaceae** | **16** | Scrophulariaceae | 4 | Amaranthaceae | 1 |
| **Plantaginaceae** | **16** | Apocynaceae | 3 | Aristolochiaceae | 1 |
| **Caprifoliaceae** | **13** | Convolvulaceae | 3 | Berberidaceae | 1 |
| **Lamiaceae** | **13** | Cystopteridaceae | 3 | Bignoniaceae | 1 |
| **Orchidaceae** | **13** | Droseraceae | 3 | Butomaceae | 1 |
| **Ranunculaceae** | **13** | Euphorbiaceae | 3 | Cornaceae | 1 |
| **Salicaceae** | **13** | Geraniaceae | 3 | Cucurbitaceae | 1 |
| **Asparagaceae** | **12** | Lentibulariaceae | 3 | Elaeagnaceae | 1 |
| Cupressaceae | 9 | Linaceae | 3 | Ginkgoaceae | 1 |
| Fagaceae | 9 | Lythraceae | 3 | Iridaceae | 1 |
| Rubiaceae | 9 | Sapindaceae | 3 | Juncaginaceae | 1 |
| Saxifragaceae | 9 | Solanaceae | 3 | Lauraceae | 1 |
| Betulaceae | 8 | Typhaceae | 3 | Menyanthaceae | 1 |
| Crassulaceae | 8 | Xanthorrhoeaceae | 3 | Myrtaceae | 1 |
| Oleaceae | 8 | Alismataceae | 2 | Nyctaginaceae | 1 |
| Amaryllidaceae | 7 | Anacardiaceae | 2 | Paulowniaceae | 1 |
| Campanulaceae | 7 | Araceae | 2 | Pittosporaceae | 1 |
| Gentianaceae | 7 | Balsaminaceae | 2 | Platanaceae | 1 |
| Onagraceae | 7 | Cannabaceae | 2 | Plumbaginaceae | 1 |
| Orobanchaceae | 7 | Cistaceae | 2 | Polygalaceae | 1 |
| Ericaceae | 6 | Ebenaceae | 2 | Rutaceae | 1 |
| Malvaceae | 6 | Hydrangeaceae | 2 | Scheuchzeriaceae | 1 |
| Polygonaceae | 6 | Hypericaceae | 2 | Smilacaceae | 1 |
| Adoxaceae | 5 | Juglandaceae | 2 | Verbenaceae | 1 |

### Appendix 2 (Figure A2)

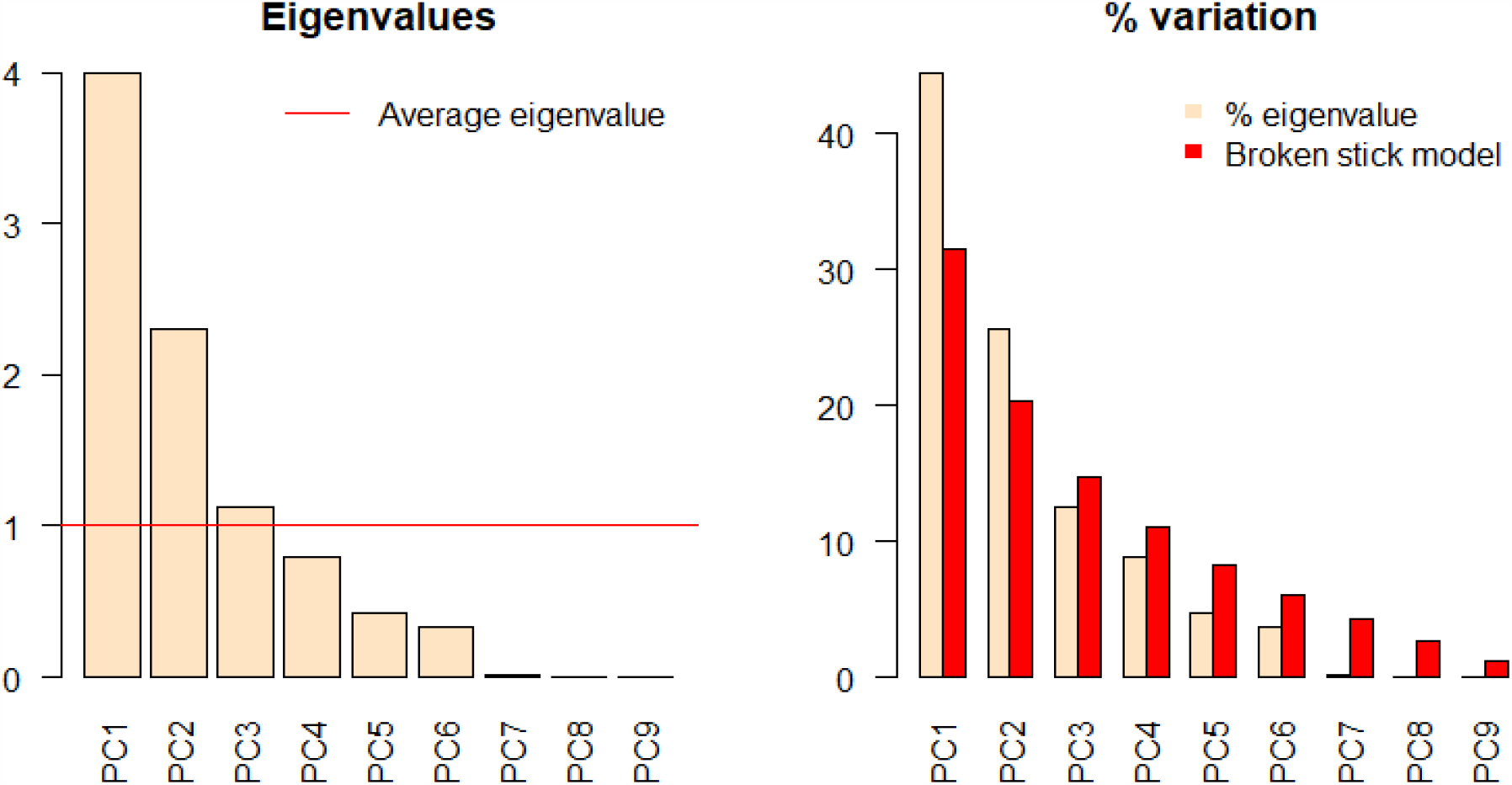
Eigenvalues of each component of the PCA (left), and their comparison with expected eigenvalues, computed following the broken stick model. Accordingly, a component is retained if its associated eigenvalue is larger than the value given by the broken stick distribution.

### Appendix 3 (Figure A3)

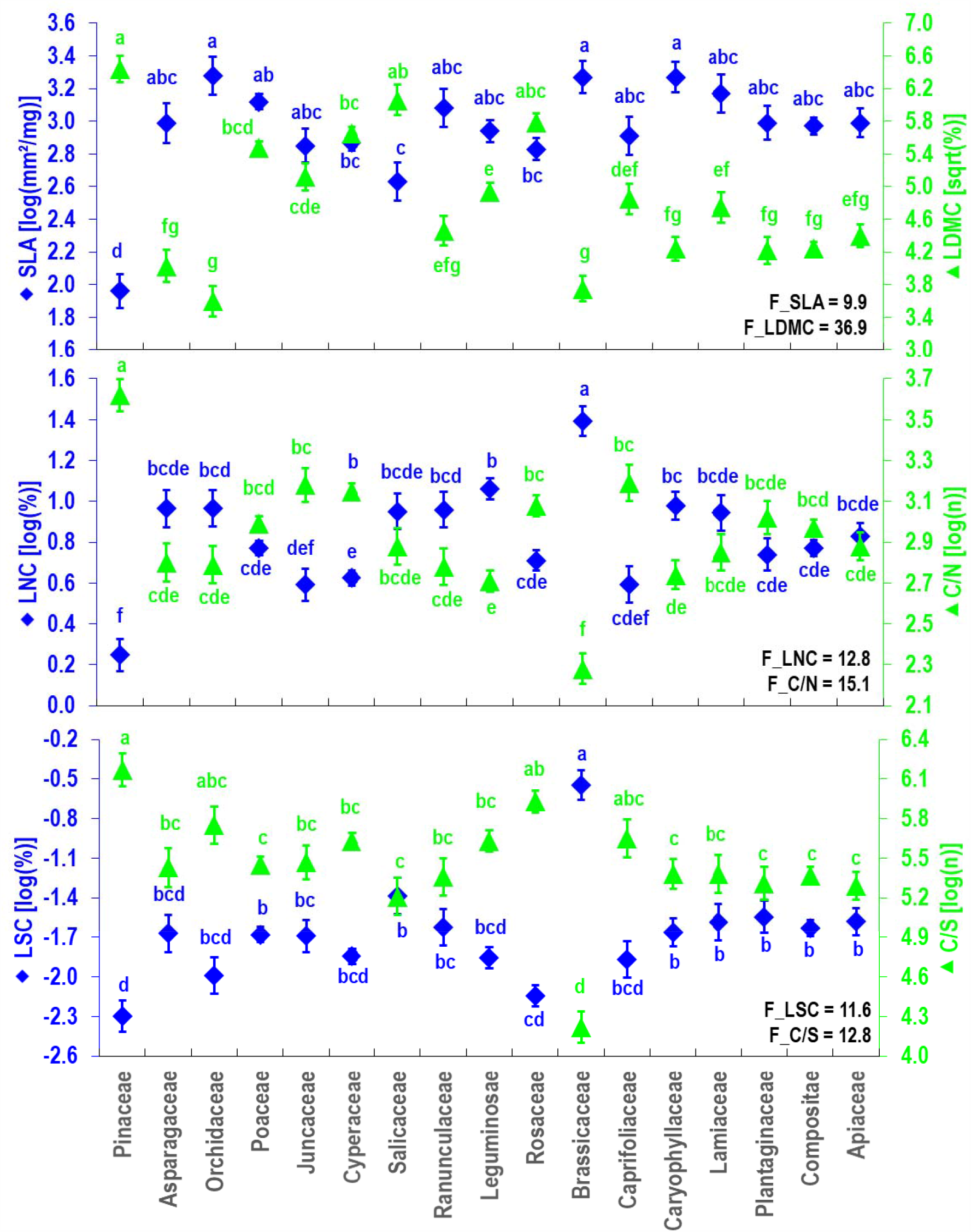
Mean values and standard error (vertical lines) of traits involved in the LES (Leaf Economics Spectrum) among plant families represented by at least 10 species. Small letters represent post-hoc comparisons. All the ANOVAs were highly statistically significant (p < 0.001, F-value reported in the figure). Legend: C/N = carbon to nitrogen ratio, C/S = carbon to sulfur ratio, LDMC = leaf dry matter content, LNC = leaf nitrogen content, LSC = leaf sulfur content, SLA = specific leaf area.

